# Obstruction of pilus retraction stimulates bacterial surface sensing

**DOI:** 10.1101/157727

**Authors:** Courtney K. Ellison, Jingbo Kan, Rebecca S. Dillard, David T. Kysela, Cheri M. Hampton, Zunlong Ke, Elizabeth R. Wright, Nicolas Biais, Ankur B. Dalia, Yves V. Brun

## Abstract

Surface association provides numerous fitness advantages to bacteria. Thus, it is critical for bacteria to recognize surface contact and to consequently initiate physiological changes required for a surface-associated lifestyle (*1*). Ubiquitous microbial appendages called pili are involved in sensing surfaces and mediating downstream surface-associated behaviors (*2*–*6*). The mechanism by which pili mediate surface sensing remains unknown, largely due to the difficulty to visualize their dynamic nature and to directly modulate their activity without genetic modification. Here, we show that *Caulobacter crescentus* pili undergo dynamic cycles of extension and retraction that cease within seconds of surface contact, and this arrest of pilus activity coincides with surface-stimulated holdfast synthesis. By physically blocking pili, we show that imposing resistance to pilus retraction is sufficient to stimulate holdfast synthesis in the absence of surface contact. Thus, resistance to type IV pilus retraction upon surface attachment is used for surface sensing.

**One Sentence Summary:** Bacteria use the tension imparted on retracting pilus fibers upon their binding to a surface for surface sensing.

## Main Text

It has been proposed that the tension imparted on retracting pili upon their attachment to a surface is used for surface sensing (*2*, *3, 5*, *6*). Here, we use the rapid surface contact-stimulated synthesis of the adhesive holdfast of *C. crescentus* to study the role of pili in surface sensing. Newborn, non-reproductive swarmer cells harbor multiple tad pili (Fig. 1A and fig. S1, A and B) and a flagellum at the same pole (fig. S2). When swarmer cells encounter a surface, holdfast synthesis at the flagellar pole is stimulated within seconds in a tad pili-dependent process (*4, 5*). Since tad pili were not known to retract, and they lack a retraction ATPase, we sought to fluorescently label them to study their behavior. Inspired by a technique for labeling flagellar filaments (*7*), we replaced a native residue within the major pilin subunit with a cysteine for subsequent labeling with thiol-reactive maleimide dyes (AF488- or AF594-mal) (Fig. 1B and fig. S3, C and E). The *C. crescentus* PilAT36C strain had wild-type levels of attachment as well as sensitivity to the pilus-dependent bacteriophage ΦCbK (fig. S4A and B), and cryo-electron tomography (cryo-ET) showed that neither the introduction of the cysteine residue nor the addition of maleimide dye affected the structure of the pilus fiber or the cell envelope (Fig. 1D and SM1). Upon fluorescence labeling, we observed tad pili and their dynamics by time-lapse microscopy (Fig 1C and MS2). The labeled pili were highly dynamic and were capable of both extension and retraction (Fig. 1C).

**Figure 1.**
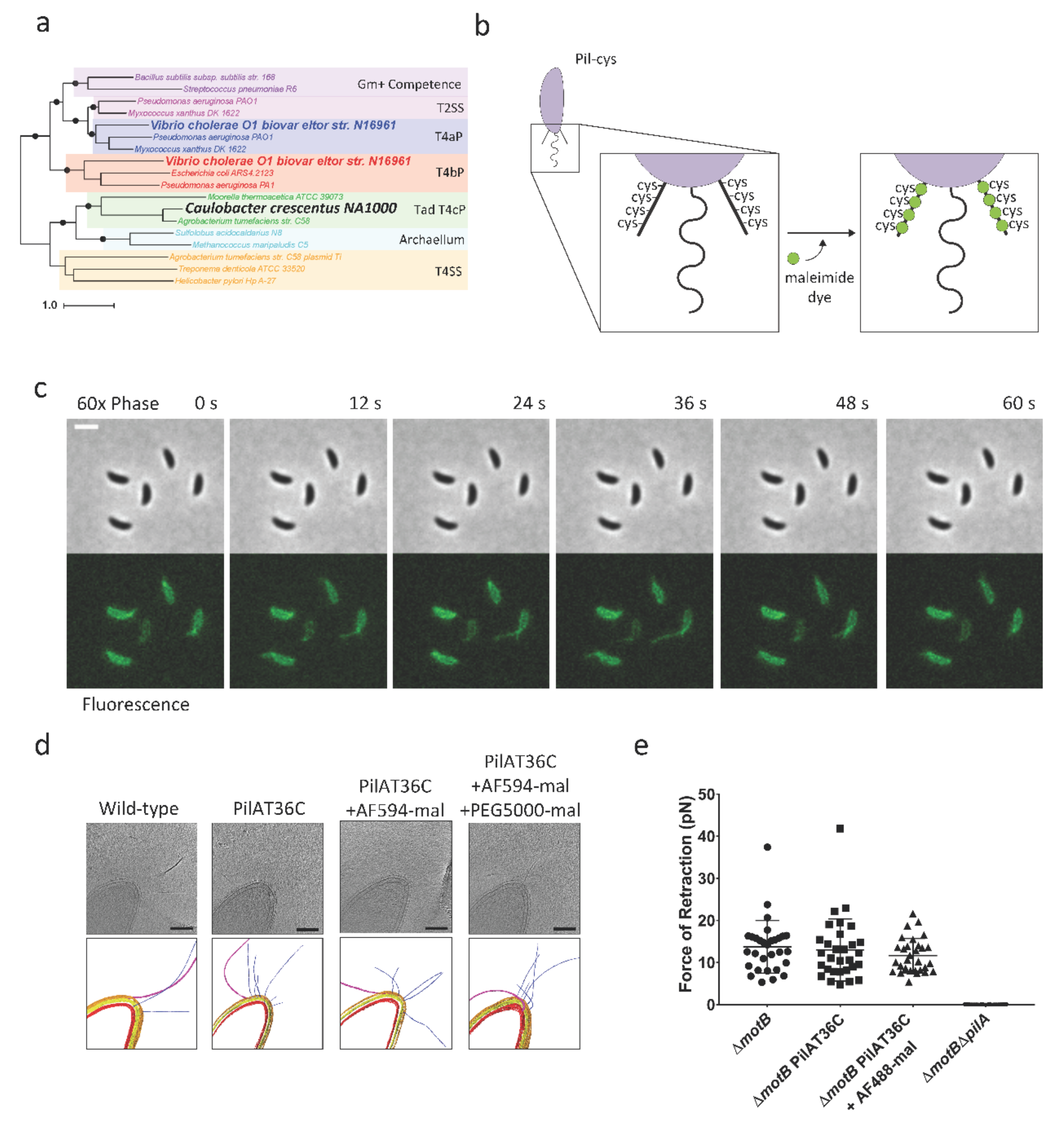
Tad type IVc pili undergo dynamic cycles of extension and retraction. (a) Phylogenetic tree comparing secretion machinery reveals a distinct tad type IVc subtype of type IV pili. An unrooted, maximum likelihood phylogeny shows relationship between type IVa (T4aP), type IVb (T4bP), IVc (T4cP) pili, archaellum (archaeal flagellum), type II secretion system (T2SS), and type IV secretion system (T4SS) extension ATPases. Circles indicate key nodes with bootstrap support values over 85%. Bolded species names correspond to pili targeted for labeling in this work. (b) Schematic of labeling pili using cysteine knock-ins. Green dots represent maleimide dye conjugates. (c) Time-lapse of labeled, synchronized Pi1AT36C swarmer cells extending and retracting pili after labeling with AF488-mal dye. Scale bar is 2 μm. (d) Slices from tomograms and corresponding 3D segmentations of wild-type, PilAT36C, PilAT36 labeled with AF594-mal, and PilAT36C blocked with PEG5000-mal and labeled with AF594-mal. In 3D segmentation volumes, flagella are pink, pili are blue, S-layer is gold, outer membrane is yellow, and inner membrane is red. Scale bars are 200 nm. (e) Micropillars assay force measurements of retraction of tad type IVc pili in flagellar motor mutant Δ*motB* strains. Flagellar motor mutants exhibiting paralyzed flagella were used to ensure all measurements obtained were dependent solely on pilus activity. Error bars show mean ± SD from 30 cells for each dataset.

Cells synthesize an average of two pili per minute at an average length of 1.08 μm (fig. S5, A and B). To measure the strength of pilus retraction, we used an established elastic micropillars assay (*8*). In this assay, pili from the same cell bind adjacent micropillars, and their subsequent retraction causes micropillar bending, which allows calculation of a retraction rate and force (fig. S6A and MS3, MS4, MS5, MS6). Based on these assays, we determined that *C. crescentus* pilus retraction generates an average force of 14 pN and that labeling does not affect the force of retraction (Fig. 1E). Extension and retraction rates of labeled pili under agar pads averaged 0.14 and 0.16 μm/sec respectively, which correlate with retraction rates measured by the micropillars assay (fig. S6, B and C). Thus, tad pili retract, and labeled tad pili extend and retract normally. Tad pili, which we refer to as type IVc (Fig. 1A and fig. S1, A and B), are more similar to the type IV secretion system and the archaellum than to other type IV pili. Labeling of the *Vibrio cholerae* type IVa MSHA (<u>m</u>annose-<u>s</u>ensitive <u>h</u>emagglutanin) pili and the *V. cholerae* type IVb TCP (<u>t</u>oxin <u>c</u>o-regulated <u>p</u>ilus) pili (fig. S3, A, B, and C and fig. S7, A and B) showed that this pili labeling method is broadly applicable to diverse pilus systems.

Direct demonstration of pilin subunit recycling during pilus synthesis has proven difficult due to limitations in techniques for studying their real-time dynamics. Upon labeling with AF488-mal, we observed that piliated swarmer cells exhibited fluorescent cell bodies, suggesting that externally labeled pilins might be internalized and recycled during pilus retraction. Using the principle of size-based exclusion by the outer membrane (OM), we labeled both wild-type and the PilAT36C strain using either the large, OM-excluded, AF488-mal (720.66 Da) dye or a small, OM-permeable, BODIPY-mal (414.22 Da) dye to quantify fluorescent cell bodies in each population (Fig. 2, A and B). The OM-permeable BODIPY-mal labeled all cell types, including stalked and predivisional cells lacking pili, from either wild-type or PilAT36C strains. In contrast, the OM-excluded AF488-mal did not label wild-type cells or cells lacking pili within the PilAT36C population unless their outer membrane was compromised (Fig. 2, B and D). Based on these results, we hypothesized that cell body staining with AF488-mal was due to pilus retraction into the periplasm. To test this hypothesis, we investigated the impact of adding a large polyethylene glycol maleimide conjugate of ~5,000 Da (PEG5000-mal). Upon co-labeling pili with PEG5000-mal and AF488-mal, labeled pili no longer exhibited dynamic activity. We also observed a drastic reduction in the number of swarmer cells with fluorescent cell bodies and a concomitant increase in the number of cells with multiple fluorescent pili (Fig. 2, C and D). Cryo-ET of cells labeled with PEG5000-mal showed no effect on pilus structure or the cell envelope (Fig. 1D). Together, these results demonstrate that AF488-mal is excluded by the OM, that externally labeled pilins are retracted into an intracellular pool of subunits, and that pilus retraction can be physically obstructed. Since newly extended pili are still labeled after removal of excess dye, these results also indicate that pilins are recycled and reused. These results extend previous work using an amine labeling dye where the strong labeling of the cell body precluded conclusions about whether the pilins could be labeled in the periplasm(*9*).

**Figure 2.**
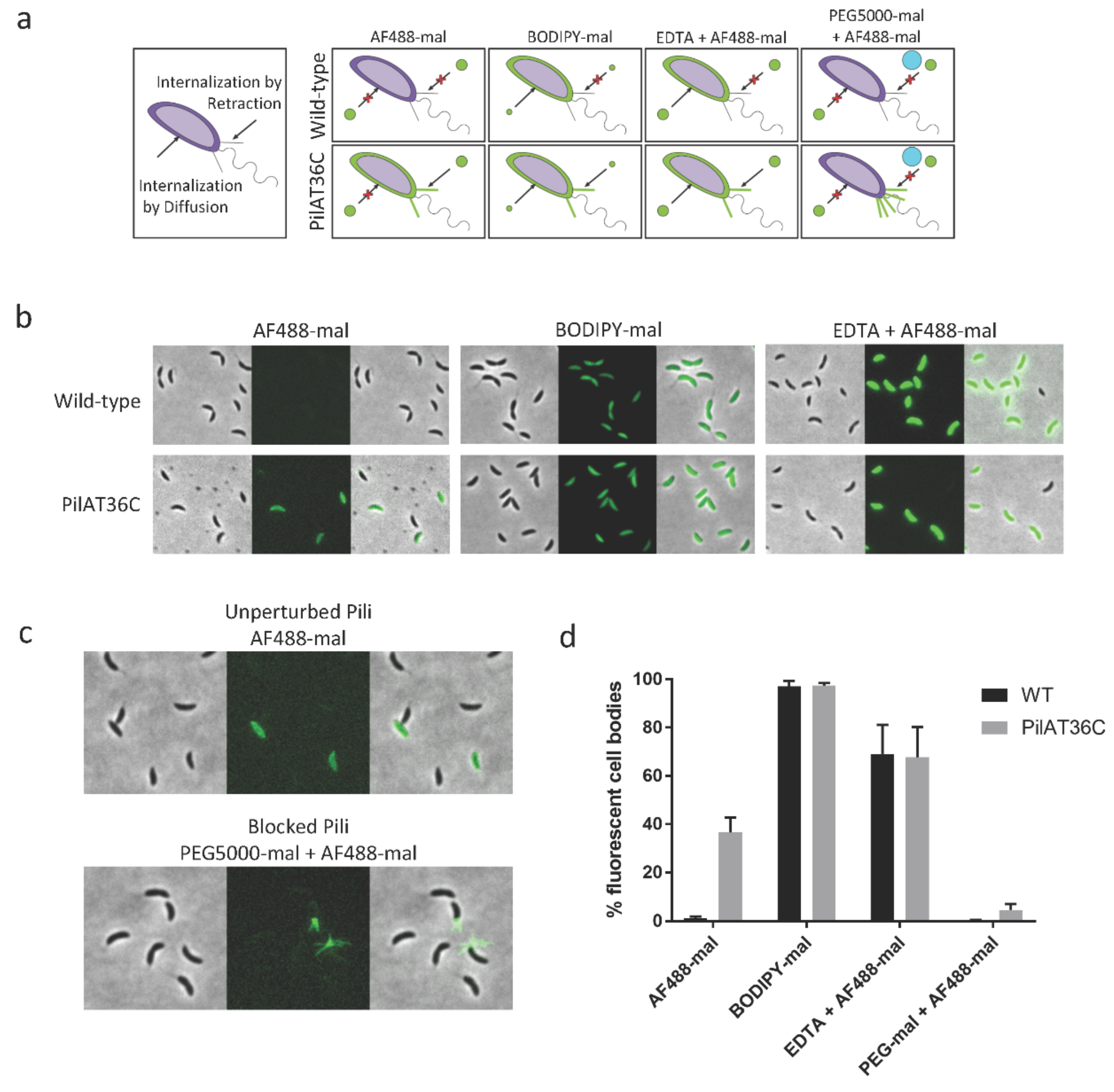
*C. crescentus* tad pilus retraction internalizes labeled pilins into a recyclable pool of subunits. (a) Labeling scheme of dye internalization for experiments shown in (b and c); green dots represent dyes of different sizes, blue ball represents PEG5000-mal, red “X” indicates inability of dye to enter cell either by diffusion or pilus retraction. (b) Representative images of wild-type or PilAT36C cells labeled with AF488-mal, BODIPY-mal, or AF488-mal after OM permeabilization with 20 mM EDTA. (c) Representative images of PilAT36C cells labeled with AF488-mal ± PEG5000-mal. (d) Quantification of fluorescent cell bodies in populations of cells from images shown in (b and c). A minimum of 398 cells from each of 3 independent biological replicates was quantified. Error bars show mean ± SD.

The inability to directly observe pilus behavior upon cell contact with a surface has complicated analysis of the role of pilus dynamics in surface sensing, requiring the use of pilus synthesis and/or retraction mutants to infer their function by observing the behavior of a nonfunctional or genetically modified system. To bypass these problems and determine the role of pilus retraction in surface sensing, we combined our abilities to observe and physically perturb fully functional pili. We first examined the dynamics of pilus extension and retraction upon surface contact. Cells with unperturbed pili arriving on the glass surface exhibited dynamic pilus activity, whereas cells blocked for pilus retraction exhibited no dynamic activity (Fig. 3, A and B and fig. S8, A and B and MS7). In unperturbed cells, we found that upon surface contact, pilus extension and retraction ceased within the same ~60 sec window as holdfast synthesis was stimulated (Fig. 3, C and D). This observation is consistent with the hypothesis that resistance on retracting surface-bound pili provides a surface-sensing signal to trigger holdfast synthesis. To test this hypothesis, we blocked pilus retraction by treatment with PEG5000-mal and measured surface attachment (Fig. 3E). Cultures in which pilus retraction was blocked experienced a 27% reduction in adhesion compared to unperturbed cells, suggesting that dynamic pilus activity is important for mediating attachment. Next, we used TIRF (total internal reflection fluorescence) microscopy to track holdfast synthesis upon surface contact in piliated swarmer cells with either unperturbed or blocked pili (Fig. 3F and fig. 9, A and B and MS8, MS9). While only 20% of unperturbed cells arrived on the surface with a holdfast synthesized prior to surface contact, 81% of cells blocked for pilus retraction had synthesized a holdfast prior to reaching the surface, indicating that blocking pilus retraction in planktonic cells was sufficient to stimulate holdfast synthesis in the absence of surface contact (Fig. 3G). Together, these data support a model in which resistance on retracting, surface-bound pili generates a surface-sensing signal.

**Figure 3.**
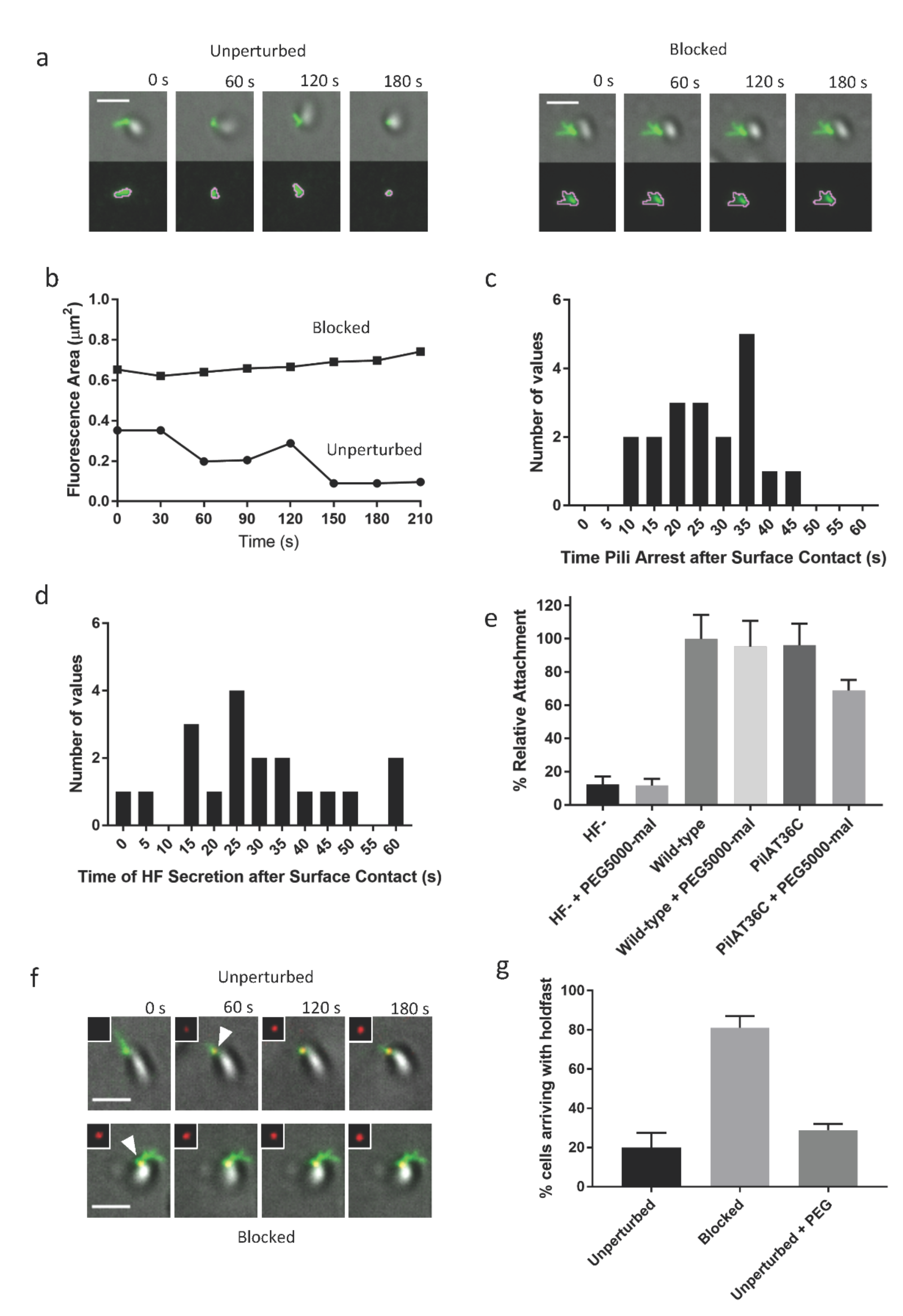
Resistance to *C. crescentus* tad pilus retraction triggers surface stimulation of holdfast synthesis. (a) Representative TIRF images of unperturbed cell labeled with AF488-mal exhibiting dynamic pilus activity and blocked cell labeled with both AF488-mal and PEG5000-mal exhibiting no dynamic pilus activity with overlays of gray cell body and green fluorescent pili. The bottom image panel shows green fluorescent pili surrounded by pink MicrobeJ overlay used to measure changes in fluorescence area of pili (μm^2^) over time shown in graph (b). (b) Graph showing changes in fluorescence area occupied by pili over time by cells shown in (a). (c) Histogram showing distribution of when dynamic pilus activity is arrested after surface contact at time = 0 s for 19 cells. (d) Histogram showing distribution of when holdfast (HF) is secreted after surface contact at time = 0 s for 19 cells. (e) Relative attachment assay showing binding efficiency of holdfast minus (HF-) and PilAT36C strains compared to wild-type after 30 min binding ± PEG5000-mal. Data are representative of binding from 3 independent cultures normalized to wild-type binding levels. Error bars show mean ± SD. PilAT36C + PEG-mal is significantly different from all other treatments based on unpaired, two-tailed T-test (*P* < 0.03). (f) Representative TIRF microscopy images of cells unperturbed or blocked for pilus retraction upon surface contact in the presence of AF594-WGA where time = 0 s is time of surface contact. The cell body is gray, labeled pili are green, and HF are red, and shown in upper right inset of each image. White arrowheads represent first appearance of holdfast for cell depicted. Scale bars are 2 μm. (g) Quantification of cells after labeling with AF488-mal (unperturbed); AF488-mal + PEG5000-mal (blocked); or AF488-mal + PEG5000 (unperturbed + PEG). A minimum of 30 cells from each of 3 independent biological replicates was quantified. Error bars show mean ± SD. Scale bars are 2 μm.

In conclusion, we have developed a broadly applicable pilus labeling method enabling real-time observation of pilus dynamics and targeted physical obstruction of this activity. We demonstrated the following, addressing several longstanding questions regarding pili and surface sensing: (1) tad pili can retract despite the absence of an orthologous retraction ATPase, (2) pilins are internalized upon retraction into a pilin pool that is recycled and reused, and (3) resistance to pilus retraction upon surface binding functions in surfacing sensing. Our method should prove valuable in determining the mechanism of tad pilus retraction and how resistance on retracting pili is sensed by the cell. Furthermore, our approach opens new avenues of research on the biology of phylogenetically related, but functionally diverse nanomachines including bacterial pili and secretion systems.

## Acknowledgements

We thank members of the Brun laboratory, Clay Fuqua, and Daniel Kearns for critical comments on the manuscript. We thank members of the Dalia laboratory Chelsea Hayes and Triana Dalia for strain construction. This work was supported by grants R01GM102841, R01GM51986, and R35GM122556 from the National Institutes of Health to YVB, by grant AI118863 from the National Institutes of Health to ABD, by National Science Foundation fellowship 1342962 to CKE, and by grant AI116566 from the National Institutes of Health to NB. This work was supported in part by grants R01GM104540 and R01GM104540-03S1 from the National Institutes of Health, NSF grant 0923395, and grants from Emory University, Children’s Healthcare of Atlanta, the Georgia Research Alliance, the Center for AIDS Research at Emory University (P30 AI050409), James B. Pendleton Charitable Trust to E.R.W. We thank the Robert P. Apkarian Integrated Electron Microscopy Core of Emory University for microscopy services and support.

## Relative Author Contributions

CE and YB designed and coordinated the overall study. CE designed and performed the *Caulobacter* experiments. DK performed the phylogenetic analysis. EW designed and RD, CH, and ZK performed the cryoET experiments. AD and CE designed and performed the *Vibrio* experiments. NB designed and JK performed the micropillar experiments. All authors analyzed and interpreted data. CE and YB wrote the manuscript with help from all authors.

